# TORC1-containing signaling endosomes source membrane from vacuoles

**DOI:** 10.1101/2024.07.02.601695

**Authors:** Kenji Muneshige, Riko Hatakeyama

**Affiliations:** Institute of Medical Sciences, School of Medicine, Medical Sciences & Nutrition, University of Aberdeen

**Author notes:** Corresponding author; Institute of Medical Sciences, University of Aberdeen, Foresterhill, Aberdeen AB25 2ZD, UK.

**Keywords:** yeast, signaling endosome, vacuole, Target of Rapamycin Complex 1 (TORC1), Atg18

## Abstract

Organelle biogenesis is fundamental to eukaryotic cell biology. Yeast signaling endosomes were recently identified as a signaling platform for the evolutionarily conserved Target of Rapamycin Complex 1 (TORC1) kinase complex. Despite the importance of signaling endosomes for TORC1-mediated control of cellular metabolism, how this organelle is generated has been a mystery. Here, we developed a system to induce synchronized *de novo* formation of signaling endosomes, enabling real-time monitoring of their biogenesis. Using this system, we identify vacuoles as a membrane source for newly formed signaling endosomes. Membrane supply from vacuoles is mediated by the CROP membrane-cutting complex, consisting of Atg18 PROPPIN and retromer subunits. The formation of signaling endosomes requires TORC1 activity, suggestive of a tightly regulated process. This study unveiled the first mechanistic principles and molecular participants of signaling endosome biogenesis.

**Summary:** Membrane is supplied by vacuoles during biogenesis of yeast signaling endosomes, in a process mediated by the membrane-cutting CROP complex and promoted by TORC1 kinase activity.

## Introduction

Eukaryotic cells are compartmentalized by membrane structures, or organelles, that perform specialized functions. Organelle biogenesis is fundamental for establishing cellular architecture and functions. Here we examine the biogenesis of yeast signaling endosomes, a novel subpopulation of endosomes (or a novel organelle; see Discussion) we recently reported (Hatakeyama & De Virgilio, 2019a).

Signaling endosomes play critical roles in the regulation of cellular metabolism. Signaling endosomes accommodate on their surface the Target of Rapamycin Complex 1 (TORC1) kinase complex, an evolutionarily conserved master regulator of cell growth and metabolism (Shimobayashi & Hall, 2014)(Liu & Sabatini, 2020). Being activated and inactivated by pro- and anti-growth stimulations, respectively, TORC1 regulates various cellular processes by phosphorylating its substrate proteins. In yeast, TORC1 localizes to the surface of two organelles: vacuoles (the yeast counterpart of lysosomes) and signaling endosomes (Sturgill et al., 2008)(Hatakeyama et al., 2019). Because TORC1 can only physically interact with and phosphorylate its nearby proteins, the difference in its localization translates into difference in its target substrates, and so the downstream biological processes controlled. We have so far reported three specific substrates of the signaling endosome-residing TORC1 pool (Hatakeyama et al., 2019)(Hatakeyama & De Virgilio, 2019b)(Chen et al., 2021): Atg13, a regulator of macroautophagy (Kamada et al., 2010); Vps27, a subunit of ESCRT-0 complex that regulates microautophagy (Oku et al., 2017); and Fab1, a lipid kinase that generates phosphatidylinositol 3,5-bisphosphate (PI(3,5)P_2_) from phosphatidylinositol 3-phosphate (PI3P) (Hasegawa et al., 2017). The vacuolar TORC1 pool, on the other hand, phosphorylates Sch9, a regulator of protein synthesis (Hatakeyama et al., 2019)(Caligaris et al., 2023). Such division of labor between spatially distinct pools of TORC1 explains, at least in part, why TORC1 can regulate diverse biological processes operating at different subcellular locations.

The principle of spatially and functionally distinct TORC1 pools seems to be evolutionarily conserved, as functional differences between lysosomal and cytoplasmic TORC1 pools were recently reported for mammalian cells (Fernandes et al., 2023). To date, however, signaling endosomes have only been described in budding yeast. Of note, the term “signaling endosome” also occurs in different contexts, either as a general concept or for other signaling pathways (Platta & Stenmark, 2011). These are distinct from the TORC1-containing signaling endosomes investigated in this study.

Apart from TORC1, signaling endosomes accommodate specific proteins and lipids that overlap with residents of vacuoles and/or canonical, TORC1-negative endosomes. The proteins shared by signaling endosomes and vacuoles include the EGO complex and Pib2, upstream regulators of TORC1 (Hatakeyama et al., 2019)(Nicastro et al., 2017)(Hatakeyama, 2021); Fab1, aforementioned lipid kinase (Chen et al., 2021); Ypt7, a yeast Rab7 small GTPase (Füllbrunn et al., 2024); Ivy1, a phospholipid-binding protein that regulates TORC1 and Fab1 (Gao et al., 2022); and Yck3, a casein kinase (Grziwa et al., 2023). Signaling endosomes also accommodate proteins typically found on endosomes but not vacuoles such as Vps21, a yeast Rab5 small GTPase (Hatakeyama et al., 2019); Vps8, a subunit of the CORVET vesicle tethering complex (Chen et al., 2021); and aforementioned Vps27 (Hatakeyama et al., 2019). The phosphoinositide composition of the signaling endosomal membrane, i.e., enrichment of PI(3,5)P_2_ and PI3P, is characteristic of both signaling endosomes and the vacuolar membrane (Chen et al., 2021). The “vacuole-endosome hybrid” composition of the signaling endosomal membrane suggests its close relation to both vacuoles and canonical endosomes. Moreover, signaling endosomes are always found adjacent to vacuoles. It is however unclear whether and how materials are exchanged between signaling endosomes and vacuoles, or signaling endosomes and canonical endosomes.

Curiously, not all cells in a population have signaling endosomes. Depending on the yeast strain background and exact growth conditions, typically, less than half of cells harbor detectable signaling endosomes at a time. Those that do usually have only one, sometimes two, signaling endosomes. These observations suggest that signaling endosomes are dynamically generated and turned over in a tightly regulated manner, rather than being a long-lived static structure. Such dynamics of signaling endosomes are likely to have significant consequences for cellular metabolism, such as autophagic activity regulated by TORC1 signaling. Currently, however, the molecular mechanisms and the upstream stimuli/triggers of signaling endosome formation are unknown. Nor do we know where signaling endosomes come from, i.e., their membrane source.

In the present work, we addressed these fundamental questions by developing a system to induce and synchronize the formation of signaling endosomes, enabling real-time observation of their biogenesis. We obtained evidence that newly formed signaling endosomes source membrane from adjacent vacuoles. Mechanistically, this process requires the Atg18 protein, and is supported by TORC1 activity. Our work thus provides a first clue for understanding signaling endosome biogenesis.

## Results

### Atg18 is required for signaling endosome formation

Studying the biogenesis of signaling endosomes in a wild-type cell population is challenging, as we cannot predict in which cells biogenesis will happen or when. To overcome this difficulty, we designed a strategy to induce and synchronize *de novo* formation of signaling endosomes. Our strategy involved two steps (Fig. 1A). Our first step was to identify a gene, “Gene X”, that is required for signaling endosome formation. The *gene XΔ* strain should have no signaling endosomes. In the second step, Gene X expression is placed under control of a conditionally inducible promotor. In such a yeast strain, without Gene X induction, cells should behave as *gene XΔ*, and thus should not have signaling endosomes. Once the expression of Gene X is induced, cells should start generating signaling endosomes, eventually behaving like wild-type cells. Time-lapse imaging during Gene X induction should enable real-time monitoring of *de novo* signaling endosome formation within a cell population.

**Figure 1.**
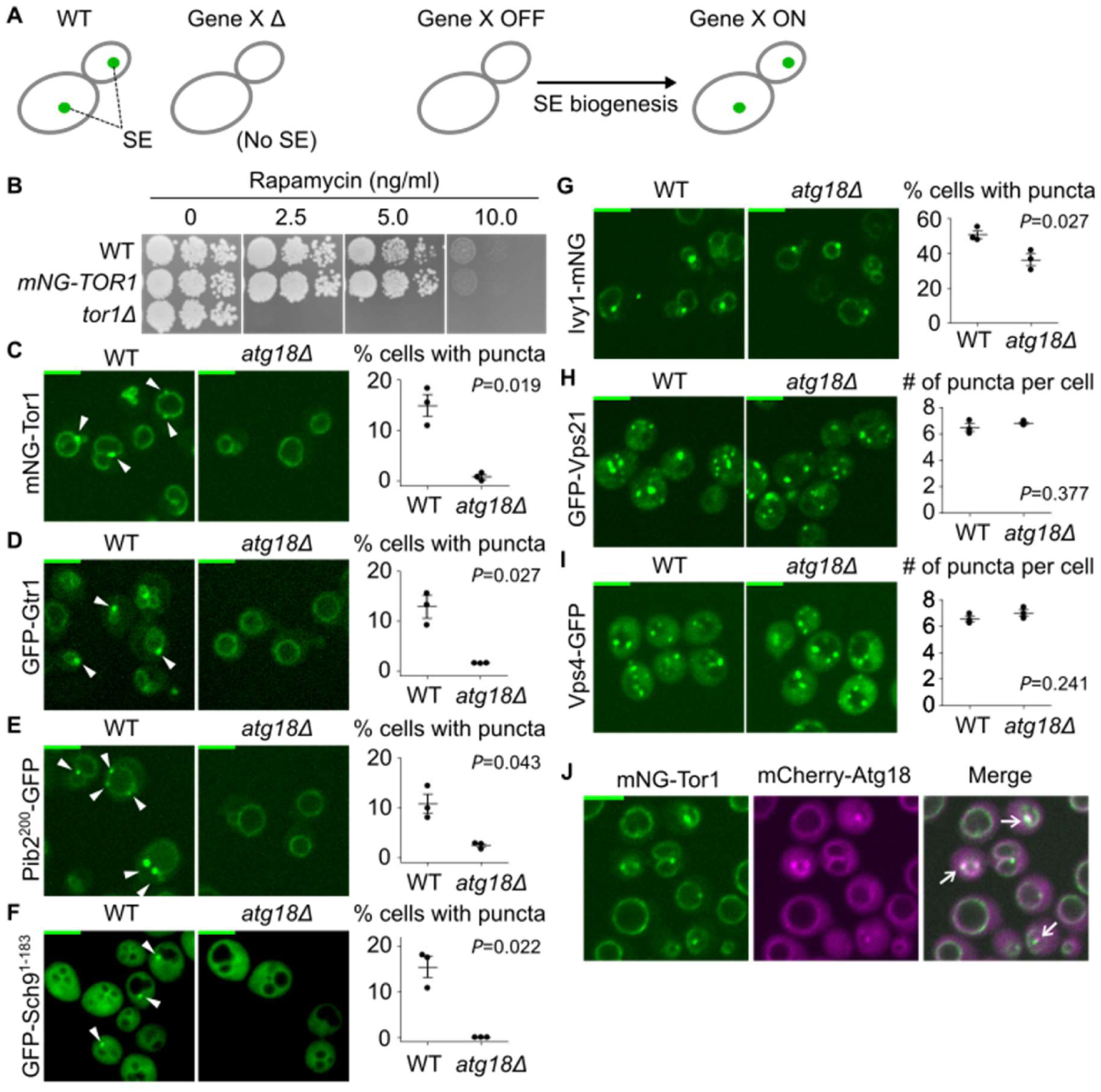
Atg18 is required for signaling endosome formation. (A) Strategy to induce and synchronize *de novo* formation of signaling endosomes (SE). See text for details. (B) Functionality of mNeonGreen-Tor1 (mNG-Tor1) used in this study. Serial dilutions of the indicated strains were spotted and grown on YPD agar plates containing rapamycin at the indicated final concentration. (C)-(I) Requirement of Atg18 for the formation of signaling endosomes but not that of canonical endosomes. Wild-type and *atg18Δ* cells expressing the indicated fusion proteins were analyzed by fluorescent microscopy. Arrowheads indicate signaling endosomes. Scale bars: 5 µm. Quantifications of puncta-containing cells are shown on the right (mean±SEM, n=3, >100 cells were analyzed for each biological replicate). In (H) and (I), because all the cells contained dots, the number of puncta per cell is presented. (J) Localization of Atg18 to signaling endosomes. Cells co-expressing mNeonGreen-Tor1 and mCherry-Atg18 were analyzed by fluorescent microscopy. Arrows indicate colocalization of Tor1 and Atg18 at signaling endosomes. Scale bar: 5 µm.

A candidate Gene X was *ATG18,* which encodes a protein of the PROPPIN (*β*-propellers that bind polyphosphoinositides) family (Michell et al., 2006). We previously reported that *atg18Δ* cells lose the perivacuolar, punctual localization of the PI(3,5)P_2_ probe GFP-Sch9^1-183^ that represents signaling endosomes (Chen et al., 2021). This observation suggested that *atg18Δ* cells may lack signaling endosomes. We tested this possibility by visualizing Tor1, a catalytic subunit of TORC1 that defines signaling endosomes. First we confirmed that mNeonGreen-tagged Tor1 is functional (Figure 1B). We found that *atg18Δ* cells lack perivacuolar puncta of mNeonGreen-Tor1 (Figure 1C). For other signaling endosome marker proteins, Gtr1 (an EGO complex subunit), Pib2, and Sch9^1-183^, puncta indicative of signaling endosomes were similarly absent (Figure 1D to 1F). The loss of puncta is not due to reduced expression of marker proteins, because their vacuolar localization was intact. Note that the vacuolar enlargement observed in Figure 1 is a known phenotype of *atg18Δ* (Dove et al., 2004).

Ivy1 has been regarded as another marker of signaling endosomes. However, we noticed that Ivy1 puncta are significantly more abundant than the puncta of Tor1, Gtr1, Pib2, and Sch9^1-183^. This observation suggests that, at least, our mNeonGreen-tagged Ivy1 construct localizes to other subcellular loci (presumably other subpopulations of endosomes) as well. Consistently, the Ivy1 puncta were reduced but not absent from *atg18Δ* cells (Figure 1G). To confirm that Atg18 is not required for the formation of other endosomal subpopulations, we examined the localization of Vps21 and Vps4, endosomal markers not specific to signaling endosomes (Hatakeyama et al., 2019)(Chen et al., 2021). These proteins indeed retained their punctual localization pattern in *atg18Δ* cells (Figure 1H and 1I), suggesting that the requirement of Atg18 is specific to signaling endosomes.

Collectively, our observations suggest an essential role for Atg18 in signaling endosome formation. We found that mCherry-Atg18 localizes to signaling endosomes (Figure 1J), consistent with direct involvement of Atg18 in signaling endosome formation and/or function.

### The Atg18-containing CROP complex mediates signaling endosome biogenesis

Having established the requirement of Atg18 for signaling endosome formation, we addressed the underlying molecular mechanisms. We first tested whether this function of Atg18 is shared by other structurally similar PROPPINs Hsv2 and Atg21 (Krick et al., 2008). Signaling endosomes were observed at normal levels in *hsv2Δ* and *atg21Δ* cells (Figure 2A), suggesting that this function is unique to Atg18 among the three yeast PROPPINs.

**Figure 2.**
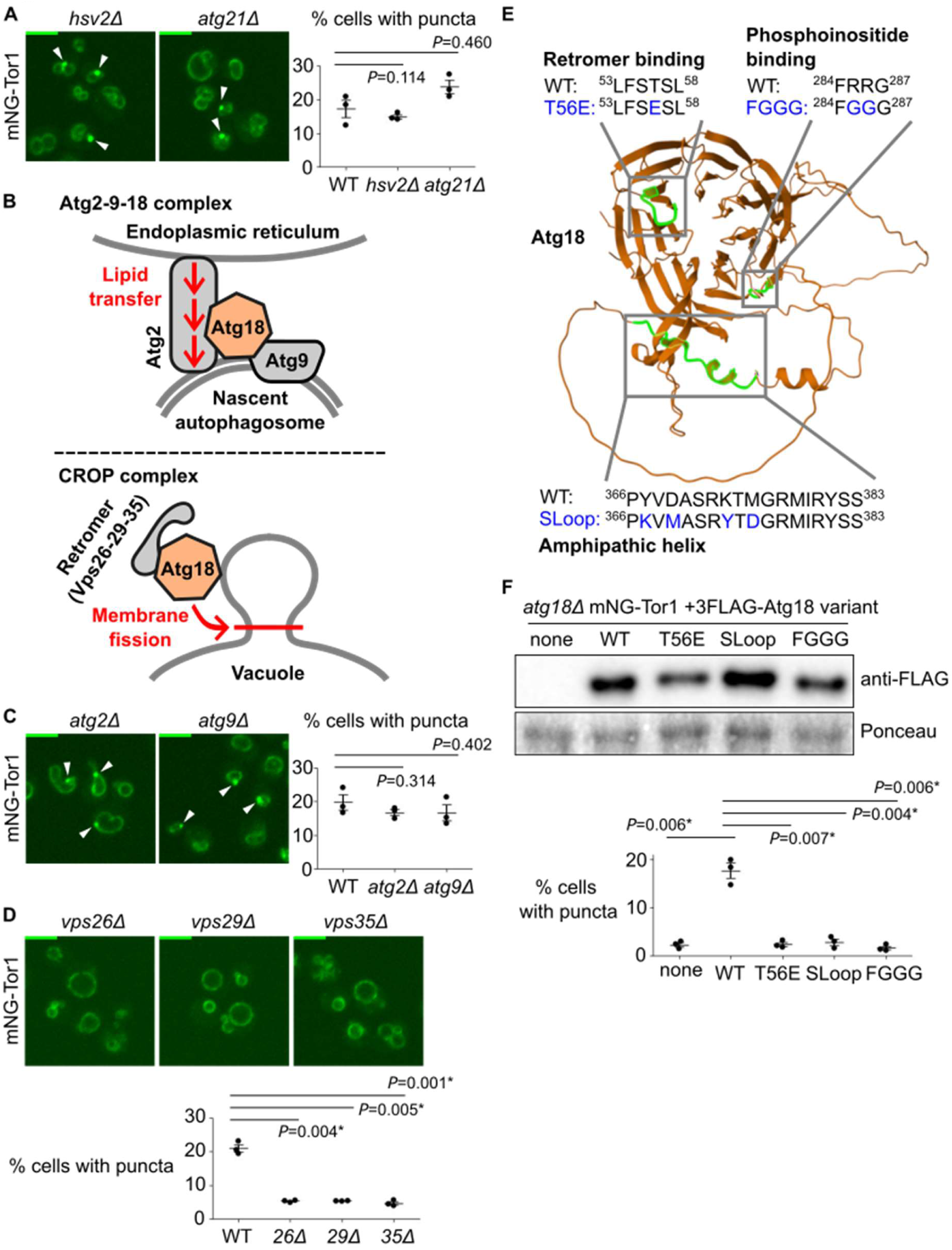
Atg18-containing CROP complex is required for signaling endosome formation. (A) No requirement of other PROPPIN proteins for signaling endosome formation. The indicated strains expressing mNeonGreen-Tor1 (mNG-Tor1) were analyzed by fluorescent microscopy. Arrowheads indicate signaling endosomes. Scale bars: 5 µm. Quantifications of puncta-containing cells are shown on the right (mean±SEM, n=3, >100 cells were analyzed for each biological replicate). (B) Compositions and functions of Atg18-containing protein complexes. See text for details. (C) No requirement of the Atg2-9-18 complex for signaling endosome formation. The indicated strains expressing mNeonGreen-Tor1 were analyzed by fluorescent microscopy. Arrowheads indicate signaling endosomes. Scale bars: 5 µm. Quantifications of puncta-containing cells are shown on the right (mean±SEM, n=3, >100 cells were analyzed for each biological replicate). (D) Requirement of the CROP complex subunits for signaling endosome formation. The indicated strains expressing mNeonGreen-Tor1 were analyzed by fluorescent microscopy. Scale bars: 5 µm. Quantifications of puncta-containing cells are shown on the right (mean±SEM, n=3, >100 cells were analyzed for each biological replicate). (E) Atg18 domains required for the formation and/or membrane-cutting activity of CROP complex. The CROP-relevant domains are highlighted in green on the AlphaFold-predicted structure of Atg18. Loss-of-function mutations examined in this study are indicated in blue. (F) Requirement of CROP-relevant Atg18 domains for signaling endosome formation. *atg18Δ* cells expressing mNeonGreen-Tor1 and the indicated FLAG-Atg18 variants were analyzed by western blotting (top) and fluorescent microscopy. Quantifications of puncta-containing cells are shown (bottom) (mean±SEM, n=3, >100 cells were analyzed for each biological replicate).

Atg18 functions in the context of distinct protein complexes (Figure 2B). The Atg18-Atg2-Atg9 trimeric complex promotes the expansion of the pre-autophagosomal membrane by transferring lipid molecules from the endoplasmic reticulum (Maeda et al., 2019)(Valverde et al., 2019)(Gómez-Sánchez et al., 2018). More recently, the Mayer and Thumm groups independently reported another Atg18-containing complex, the CROP (Cutting Retromer-on-PROPPIN) complex (Courtellemont et al., 2022)(Marquardt et al., 2023)(Gopaldass & Mayer, 2024)(Marquardt & Thumm, 2023). CROP is comprised of Atg18, Vps26, Vps29, and Vps35, the latter three of which are subunits of the retromer sorting complex (Seaman et al., 1998). CROP cuts the membrane of endo-lysosomal compartments, promoting formation of tubulo-vesicular transport carriers from endosomes, and vacuolar fission and fragmentation (De Leo et al., 2021)(Gopaldass et al., 2017)(Courtellemont et al., 2022)(Marquardt et al., 2023). We addressed the protein complex and functions through which Atg18 supports signaling endosome formation by deleting each binding partner. Signaling endosomes were intact in *atg2Δ* and *atg9Δ* cells (Figure 2C), ruling out involvement of the Atg18-Atg2-Atg9 complex. In contrast, *vps26Δ, vps29Δ* and *vps35Δ* mutants had significantly fewer signaling endosomes than wild-type cells (Figure 2D), suggesting a role for CROP-mediated membrane fission in signaling endosome formation. To corroborate this notion, we analyzed three Atg18 mutants defective in CROP complex formation and/or membrane fission, T56E, SLoop, and FGGG mutants (Figure 2E) (Courtellemont et al., 2022)(Gopaldass & Mayer, 2024). As expected, these Atg18 mutants (which were expressed well; Figure 2F top) failed to support signaling endosome formation (Figure 2F bottom).

Our results suggest a major role for the CROP complex in signaling endosome formation. We nevertheless noticed that the defects in retromer mutants (Figure 2D) are significant but only partial, differing from the complete loss of signaling endosomes in *atg18Δ* cells. This observation aligns well with the fact that Atg18 can act alone to cut a membrane, albeit less efficiently than when it is in the CROP complex (Gopaldass et al., 2017)(Courtellemont et al., 2022).

### Development of an inducible system to study signaling endosome biogenesis

The participation of the CROP complex indicates that signaling endosome biogenesis involves a membrane fission process. Because 1) the CROP complex cuts the vacuolar membrane, 2) signaling endosomes are always adjacent to vacuoles, and 3) the protein and lipid composition significantly overlap between signaling endosomes and vacuoles, it was plausible that vacuoles supply the membrane, through a fission process, to newly born signaling endosomes.

We sought to test this hypothesis by monitoring signaling endosome formation in real-time. To this end, we took advantage of the complete loss of signaling endosomes in *atg18Δ* cells, identifying *ATG18* as Gene X (Figure 1A). To conditionally express Atg18, we utilized the WTC_846_ toolkit, an improved tetracycline-inducible gene expression system (Azizoğlu et al., 2021). We confirmed that anhydrotetracycline induces FLAG-Atg18 expression in a dose-dependent manner in the *WTC_846_-ATG18* strain (Figure 3A). Without anhydrotetracycline treatment, Atg18 expression was undetectable. The Atg18 expression level with 1 ng/ml anhydrotetracycline approximately corresponded to the endogenous level (first lane), so we used this concentration of anhydrotetracycline in the following experiments unless specified otherwise. The expression level of Tor1 did not change after anhydrotetracycline treatment, meaning that anhydrotetracycline-dependent changes in Tor1 distribution described below are not due to changes in the Tor1 abundance.

**Figure 3.**
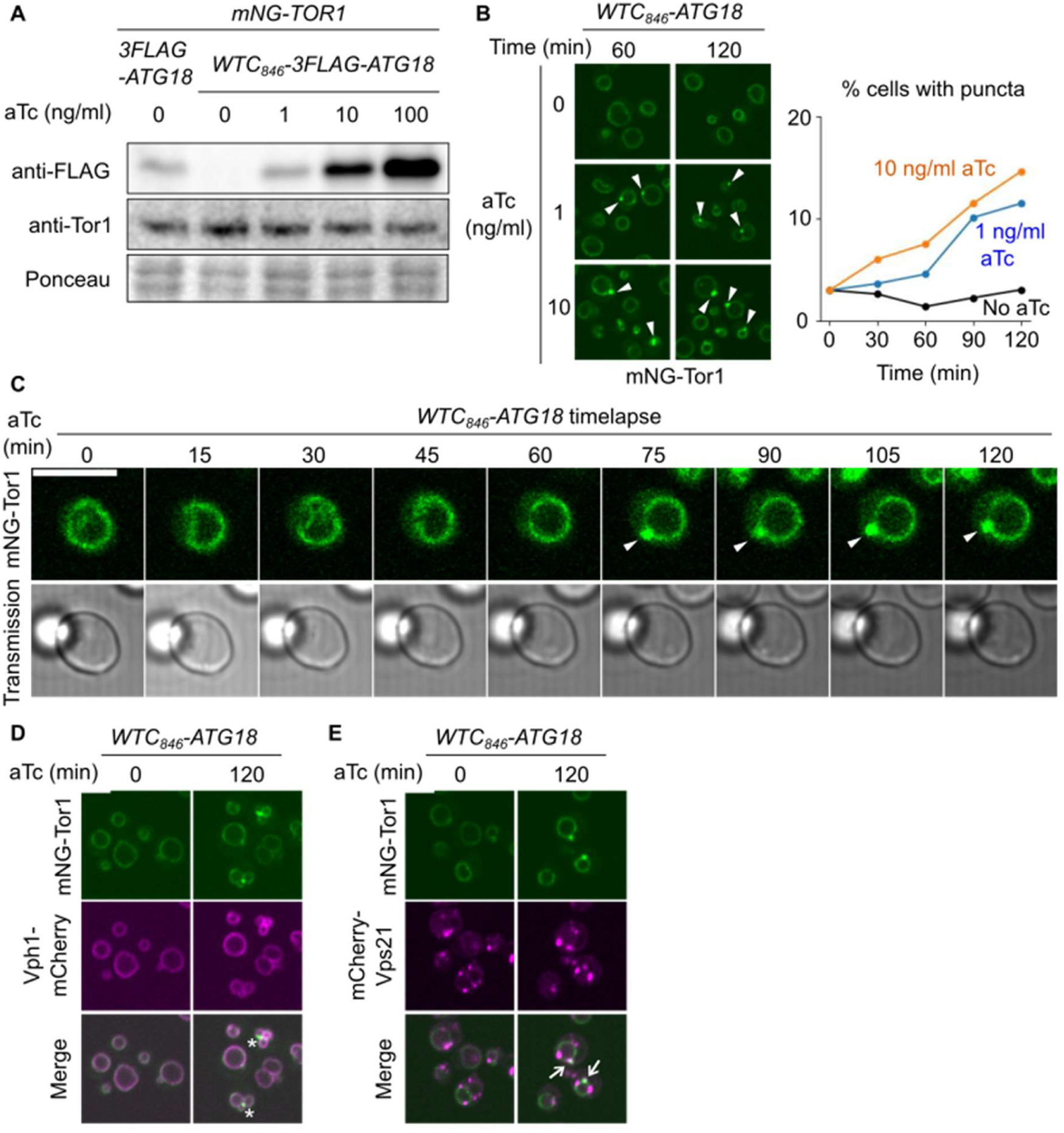
Development of a signaling endosome induction system. (A) Confirmation of WTC_846_-induced Atg18 expression. The indicated strains were treated with anhydrotetracycline (aTc) at the indicated final concentrations for 60 minutes and analyzed by western blotting. (B) WTC_846_-induced formation of signaling endosomes. *WTC_846_-ATG18* cells expressing mNeonGreen-Tor1 (mNG-Tor1) were treated with anhydrotetracycline at the indicated final concentrations for the indicated periods and analyzed by fluorescent microscopy. Arrowheads indicate newly formed signaling endosomes. Scale bar: 5 µm. Quantifications of puncta-containing cells are shown on the right (>100 cells were analyzed). (C) Time-lapse monitoring of signaling endosomes formation. *WTC_846_-ATG18* cells expressing mNeonGreen-Tor1 (mNG-Tor1) were treated with 1 ng/ml anhydrotetracycline and analyzed by time-lapse fluorescent microscopy. Arrowheads indicate newly formed signaling endosomes. Scale bar: 5 µm. (D) Absence of v-ATPase on induced signaling endosomes. *WTC_846_-ATG18* cells co-expressing mNeonGreen-Tor1 and Vph1-mCherry were treated with 1 ng/ml anhydrotetracycline for the indicated periods and analyzed by fluorescent microscopy. Asterisks indicate the absence of Vph1 puncta on newly formed signaling endosomes. Scale bar: 5 µm. (E) Localization of Vps21 to induced signaling endosomes. *WTC_846_-ATG18* cells co-expressing mNeonGreen-Tor1 and mCherry-Vps21 were treated with 1 ng/ml anhydrotetracycline for the indicated periods and analyzed by fluorescent microscopy. Arrows indicate colocalization of Tor1 and Vps21 at newly formed signaling endosomes. Scale bar: 5 µm.

As expected, *WTC_846_-ATG18* cells showed no Tor1 puncta without anhydrotetracycline treatment (Figure 3B). Tor1 puncta started to appear around 30 to 60 minutes after the addition of anhydrotetracycline. The number of Tor1 puncta correlated with the concentration of anhydrotetracycline, i.e., the expression level of Atg18. Interestingly however, not all cells formed Tor1 puncta even when Atg18 was overexpressed (10 ng/ml aTc; see Discussion).

Timelapse imaging allowed us to track specific cells and directly observe the formation of Tor1 puncta (Figure 3C). To confirm that these Tor1 puncta are *bona fide* signaling endosomes rather than fragmented vacuoles, we simultaneously visualized proteins that discriminate the two structures. We first examined the vacuolar ATPase subunit Vph1, which is present on vacuoles but not on signaling endosomes. As expected, Vph1 did not form puncta upon Atg18 expression (Figure 3D). We then examined the Rab5 small GTPase Vps21, which is present on signaling endosomes but not on vacuoles. A fraction of Vps21 indeed colocalized with Tor1 puncta (Figure 3E), further confirming the identity of these Tor1 puncta as signaling endosomes. Hence, our *WTC_846_-ATG18* system enabled direct observation of signaling endosome biogenesis.

### Signaling endosomes inherit the vacuolar membrane

Using the *WTC_846_-ATG18* system, we tested the vacuole-origin hypothesis by examining the vacuolar membrane during *de novo* formation of signaling endosomes. As a first approach, we tracked vacuolar lipids using the lipophilic dye FM4-64, an established marker of the vacuolar membrane. FM4-64, once added to the culture media, is initially inserted into the plasma membrane and then unidirectionally trafficked to vacuoles through endocytosis (Vida & Emr, 1995). When FM4-64 is washed out shortly after its addition, and allowed to complete its endocytic trafficking (which takes 30 to 60 minutes), it specifically stains the vacuolar membrane. We stained the vacuolar membrane of *WTC_846_-ATG18* cells with FM4-64 prior to Atg18 induction (Figure 4A, time 0). Because FM4-64 has been washed out, any FM4-64 signal observed afterward must have come from the vacuolar membrane. Around 30 to 60 minutes after Atg18 induction, we started observing the formation of perivacuolar, FM4-64-positive puncta. These puncta were positive for Tor1, identifying them as signaling endosomes. This observation supports the idea that newly formed signaling endosomes acquire lipids from vacuoles.

**Figure 4.**
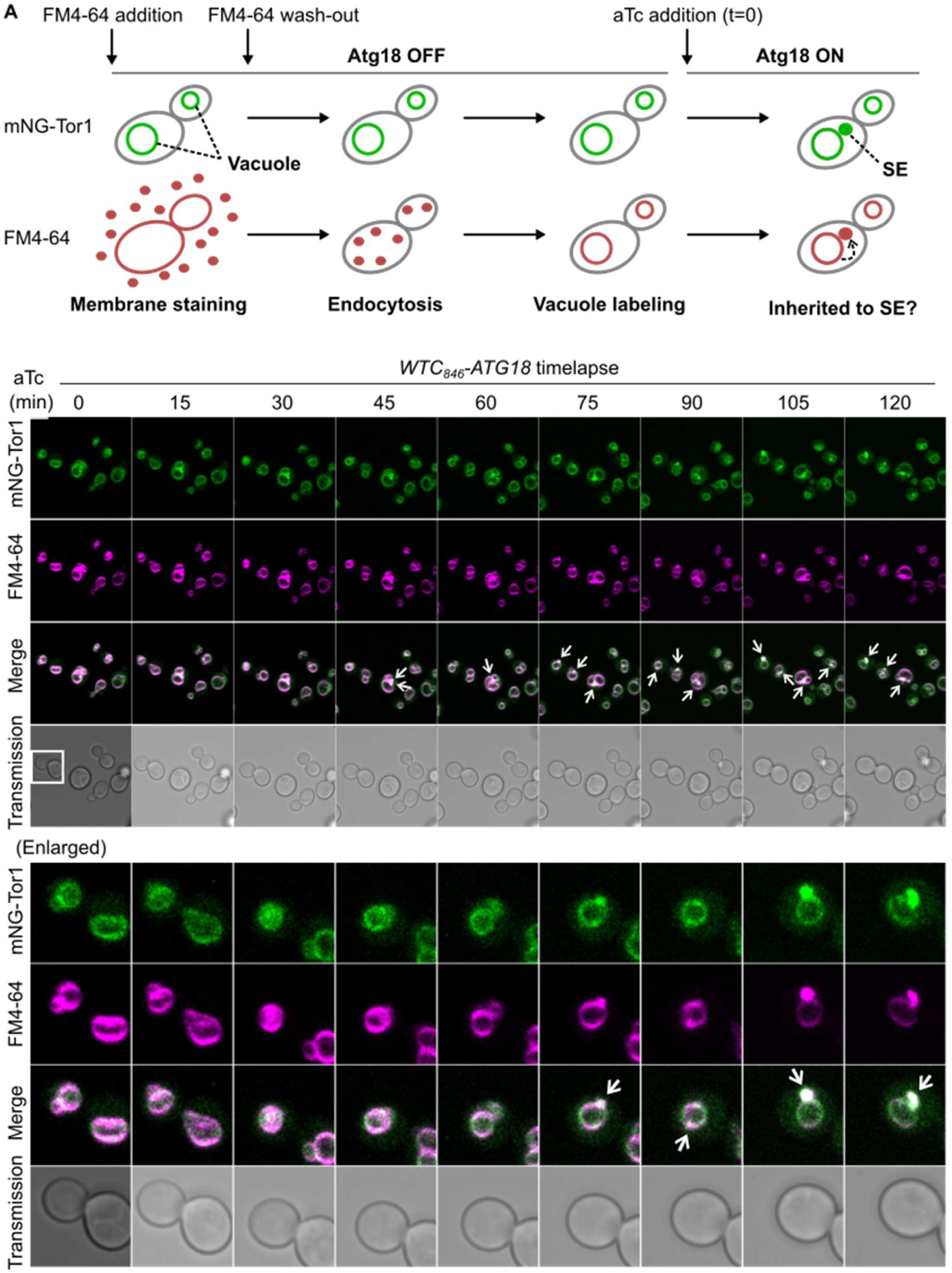
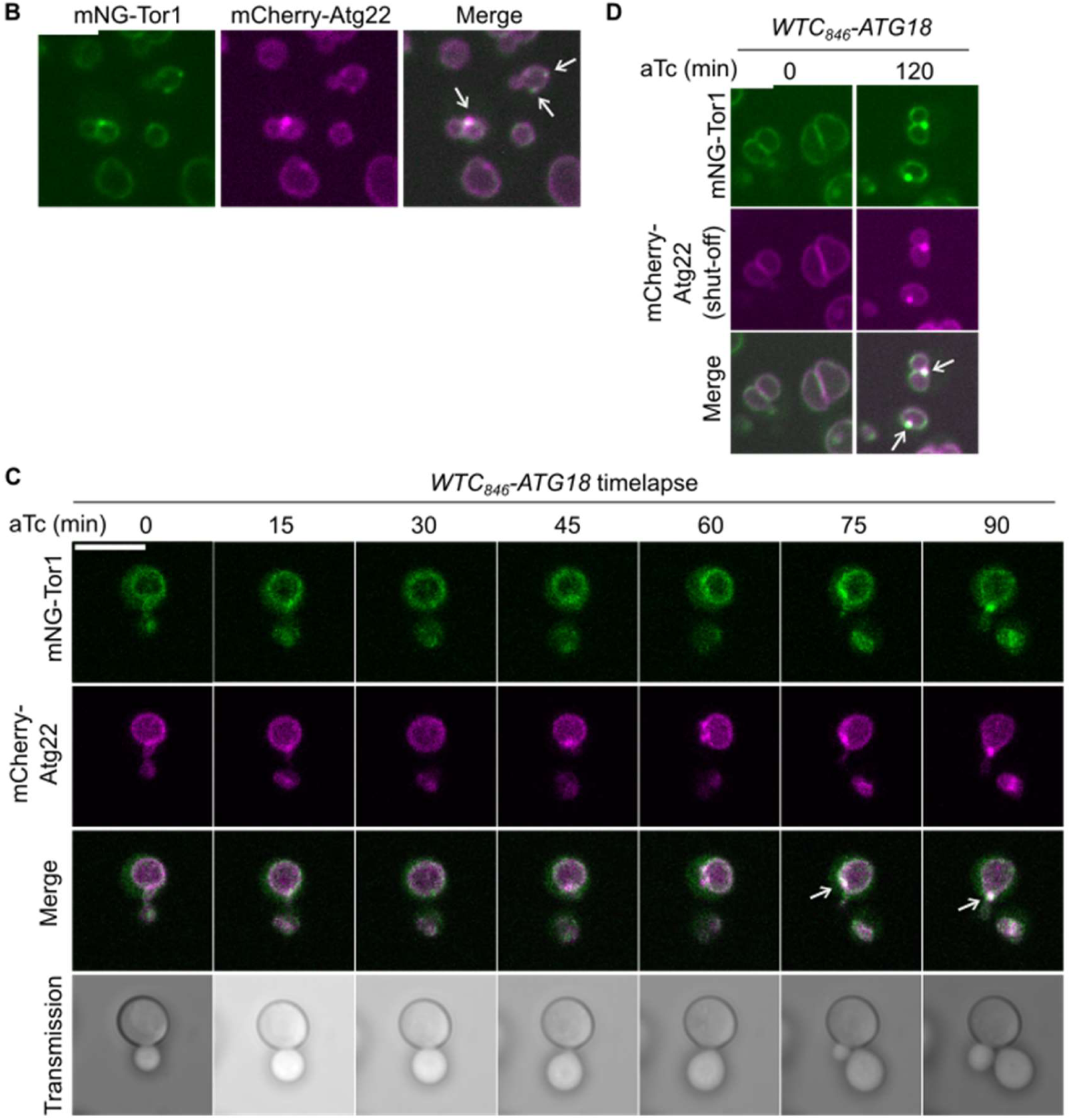
Membrane inheritance from vacuoles to newly formed signaling endosomes. (A) Inheritance of vacuolar membrane lipids to newly formed signaling endosomes. (Top) Experimental scheme. SE: signaling endosome. (Middle) *WTC_846_-ATG18* cells expressing mNeonGreen-Tor1 (mNG-Tor1) were pre-stained with FM4-64, treated with 1 ng/ml anhydrotetracycline (aTc), and analyzed by time-lapse fluorescent microscopy. Arrows indicate Tor1 localization and FM4-64 staining overlapping at newly formed signaling endosomes. Scale bar: 5 µm. Enlarged images of the cell marked by the box are shown at the bottom. (B) Localization of Atg22 to signaling endosomes. Cells co-expressing mNeonGreen-Tor1 and mCherry-Atg22 were analyzed by fluorescent microscopy. Arrows indicate colocalization of Tor1 and Atg22 at signaling endosomes. Scale bar: 5 µm. (C) Inheritance of Atg22 from the vacuolar membrane to newly formed signaling endosomes. *WTC_846_-ATG18* cells co-expressing mNeonGreen-Tor1 and mCherry-Atg22 were treated with 1 ng/ml anhydrotetracycline and analyzed by time-lapse fluorescent microscopy. Arrows indicate colocalization of Tor1 and Atg22 at newly formed signaling endosomes. Scale bar: 5 µm. (D) Inheritance of pre-existing Atg22 from the vacuolar membrane to newly formed signaling endosomes. *WTC_846_-ATG18* cells co-expressing mNeonGreen-Tor1 (under the native promotor) and mCherry-Atg22 (under the inducible *GAL* promotor) were cultured first in raffinose media and then in galactose media to induce the expression of mCherry-Atg22. Cells were then cultured in glucose media to shut off the *de novo* expression of mCherry-Atg22, treated with 1 ng/ml anhydrotetracycline for the indicated periods, and analyzed by fluorescent microscopy. Arrows indicate colocalization of Tor1 and Atg22 at newly formed signaling endosomes. Scale bar: 5 µm.

We then asked whether lipids are transferred as individual molecules, for example via lipid transfer across a membrane contact site (Scorrano et al., 2019), or in the form of a membrane. The latter scenario would be supported if lipids accompany membrane proteins. None of the known signaling endosomal proteins however serves as a suitable marker, because they are actually soluble proteins that are either peripherally attached or merely anchored to the membrane, and can potentially transfer between organelles through cytoplasmic diffusion. We therefore searched for a membrane-spanning protein that localizes to both vacuoles and signaling endosomes. Searching a genome-wide protein localization database, YeastRGB (Dubreuil et al., 2019), identified the amino acid transporter Atg22 (Yang et al., 2006) as a membrane-spanning protein localizing to vacuoles and perivacuolar puncta. We confirmed these perivacuolar puncta are signaling endosomes by colocalizing them with Tor1 puncta (Figure 4B). Atg22 is therefore the first transmembrane protein that localizes to signaling endosomes, providing a useful marker for tracking the membrane.

We tracked Atg22 during Atg18 induction. Without induction, Atg22 localized solely to the vacuolar membrane as expected (Figure 4C). Upon induction, Atg22 migrated to newly formed signaling endosomes marked by Tor1. We obtained the same result even when the expression of Atg22 was shut off (using the conditional *GAL* promotor) prior to Atg18 induction (Figure 4D), suggesting the migration of pre-existing (rather than newly synthesized) Atg22 from vacuoles to signaling endosomes.

Together, our FM4-64- and Atg22-tracking experiments demonstrate that lipids and a membrane-integrated protein migrate from the vacuolar membrane during the formation of signaling endosomes. These results establish vacuoles as a membrane source of newly formed signaling endosomes. It is however worth noting that not all the vacuolar membrane proteins are transferred to signaling endosomes. Vph1, for example, remains confined to the vacuolar membrane as already mentioned (Figure 3D; see Discussion for potential explanation).

### TORC1-Fab1 signaling promotes signaling endosome biogenesis

We next asked for the signals that trigger the formation of signaling endosomes. Our Atg18 induction system enabled us to specifically test the effect of TORC1 activity. When TORC1 was inhibited by rapamycin, Atg18-driven formation of Tor1 puncta was severely impaired (Figure 5A). We noticed a marginal impairment of Atg18 protein induction under rapamycin treatment (Figure 5B), as expected from TORC1’s role in global protein translation. This reduction however does not explain the observed lack of signaling endosome formation, because the Atg18 level induced in the presence of rapamycin was similar to its endogenous expression level (first lane).

**Figure 5.**
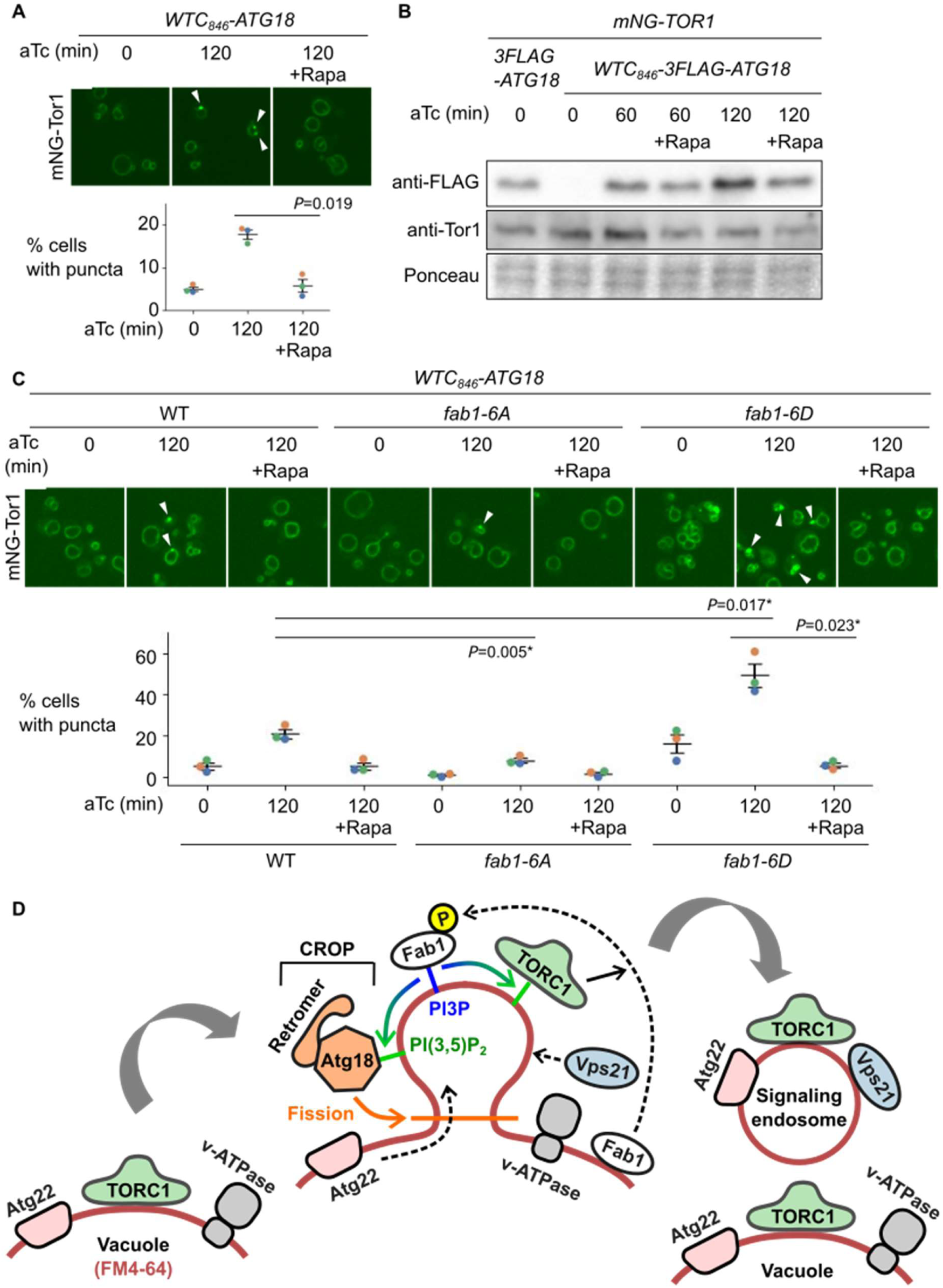
TORC1-Fab1 signaling promotes signaling endosome biogenesis. (A) Requirement of TORC1 activity for signaling endosome formation. *WTC_846_-ATG18* strains expressing mNeonGreen-Tor1 (mNG-Tor1) were treated with 1 ng/ml anhydrotetracycline (aTc) for the indicated periods in the presence or absence of 200 ng/ml rapamycin and analyzed by fluorescent microscopy. Arrowheads indicate newly formed signaling endosomes. Scale bar: 5 µm. Quantifications of puncta-containing cells are shown at the bottom (mean±SEM, n=3, >100 cells were analyzed for each biological replicate; data points are color-coded according to biological replicates). (B) Confirmation of Atg18 induction in the presence of rapamycin. The indicated strains were treated with 1 ng/ml anhydrotetracycline for the indicated periods in the presence or absence of 200 ng/ml rapamycin and analyzed by western blotting. (C) Fab1 phosphorylation by TORC1 promotes signaling endosome formation. Wild-type, *fab1-6A*, or *fab1-6D* cells expressing mNeonGreen-Tor1 were treated with 1 ng/ml anhydrotetracycline for the indicated periods in the presence or absence of 200 ng/ml rapamycin and analyzed by fluorescent microscopy. Arrowheads indicate newly formed signaling endosomes. Scale bar: 5 µm. Quantifications of puncta-containing cells are shown at the bottom (mean±SEM, n=3, >100 cells were analyzed for each biological replicate; data points are color-coded according to biological replicates). (D) Model of signaling endosome biogenesis. Fab1 phosphorylation by TORC1 triggers its relocation to the site of signaling endosome formation on the vacuolar membrane. There, Fab1 locally produces PI(3,5)P_2_ from PI3P. PI(3,5)P_2_ anchors and stabilizes TORC1, which in turn further phosphorylates and stabilizes Fab1, resulting in a positive feedback loop. In the meantime, PI(3,5)P_2_ recruits and activates the Atg18-containing CROP complex, which cuts the vacuolar membrane and thereby separates signaling endosomes. During this process, vacuolar membrane lipids (labelled by FM4-64) as well as certain membrane-spanning proteins such as Atg22 migrate into forming signaling endosomes. Other membrane-spanning proteins such as v-ATPase remain on the vacuolar membrane. Certain endosomal proteins such as Vps21 small GTPase are delivered to the membrane of newly formed signaling endosomes, completing the signaling endosome biogenesis.

Having established a requirement of TORC1 activity for signaling endosome formation, we searched for the relevant phosphorylation substrate. One TORC1 substrate functionally linked to Atg18 is the Fab1 lipid kinase, which produces PI(3,5)P_2_ that binds to Atg18 and promotes its membrane-fission activity (Gopaldass et al., 2017). TORC1 directly phosphorylates Fab1, stimulating its relocation from vacuoles to signaling endosomes (Chen et al., 2021). This triggers the local production of PI(3,5)P_2_ at signaling endosomes, which in turn anchors TORC1 to the membrane, serving as a positive feedback mechanism to maintain functional signaling endosomes. It has been unknown whether Fab1 phosphorylation also plays a role in the initial formation of signaling endosomes.

We examined the effect of Fab1 phosphorylation using its phospho-deficient (*fab1-6A*) and -mimetic (*fab1-6D*) mutants (Chen et al., 2021). Upon Atg18 induction, *fab1-6A* formed fewer signaling endosomes compared to wild-type cells (Figure 5C). Conversely, *fab1-6D* mutants formed more signalling endosomes. These observations suggest that Fab1 phosphorylation indeed promotes signaling endosome biogenesis. Rapamycin exerted its inhibitory effect even in *fab1-6D* cells, suggesting the existence of additional TORC1 substrate(s), and/or additional TORC1-phosphorylation site(s) on Fab1 that are required for signaling endosome formation.

Our results suggest a positive regulatory role for the TORC1-Fab1 signaling in *de novo* formation of signaling endosomes, presumably through recruitment and/or activation of Atg18. Given that Atg18 regulates Fab1 (Efe et al., 2007)(Jin et al., 2008), these results implicate Atg18 with the TORC1-Fab1 module in forming an extended, tightly interconnected regulatory feedback loop (Figure 5D).

## Discussion

The TORC1-positive perivacuolar structure has been designated the signaling endosome, because it accommodates proteins typically found on endosomes (Hatakeyama & De Virgilio, 2019a). In the present study, we identify vacuoles as a membrane source for newly formed signaling endosomes. This membrane source for signaling endosomes differs starkly from our current knowledge of canonical endosomes, for which the membrane is supplied mainly from the plasma membrane and the Golgi apparatus via intracellular vesicle trafficking. Our findings are somewhat unexpected and counterintuitive given the bulk membrane flow from canonical endosomes to vacuoles following endocytosis, which contrasts with the flow from vacuoles to signaling endosomes that we discovered here. This unique feature suggests that signaling endosomes should be most correctly viewed as a distinct organelle rather than a mere subpopulation of endosomes.

At the mechanistic level, the requirement for Atg18, particularly in the context of the CROP complex, strongly suggests that signaling endosomes are formed through fission of the vacuolar membrane (Figure 5D). This process would be topologically similar to the vacuolar fragmentation process, which is also mediated by the CROP complex. The two processes are however distinct in terms of symmetry. Vacuolar fragmentation is a symmetric process via which a parental vacuole is divided into qualitatively identical daughter vacuoles. In contrast, signaling endosomes differ from vacuoles in terms of composition and function. This fundamental difference between the two processes suggests the existence of molecular machinery uniquely involved in the biogenesis signaling endosome biogenesis.

Questions remain concerning how the membrane of signaling endosomes is differentiated, and maintained distinct, from the vacuolar membrane. There must exist a selection mechanism that targets certain vacuolar proteins (such as Atg22) to signaling endosomes, while retaining others (such as v-ATPase) on the vacuolar membrane (Figure 5D). One possibility is that Atg22 contains a signaling endosome-targeting motif, and/or that v-ATPase has a vacuole-retaining motif. Another possibility is that distinct microdomains of the vacuolar membrane encode the fates of resident proteins. During prolonged nutrient starvation, two distinct microdomains of the vacuolar membrane become visible, one enriched with, and the other depleted of, sterols (Toulmay & Prinz, 2013)(Kim & Budin, 2024). Interestingly, the sterol-enriched raft-like microdomain accommodates proteins including the EGO complex and Ivy1, similar to signaling endosomes. On the other hand, the v-ATPase resides in the sterol-poor microdomain. If in unstarved cells these microdomains also exist (albeit not readily visible), or are temporarily/locally formed during signaling endosome biogenesis, then signaling endosomes might originate from the sterol-rich domain, resulting in the selective inheritance of vacuolar proteins residing there.

An equally important question is how signaling endosomes acquire endosomal proteins such as the Vps21 GTPase (Figure 3E). Its stable association with the signaling endosomal membrane indicates the prior arrival of its upstream regulators such as guanine nucleotide exchange factors. How they are delivered is however a mystery. Future studies could determine whether these proteins are individually delivered to signaling endosomes or arrive by membrane fusion. The latter scenario would be consistent with the recently proposed model that a fraction of canonical endosomes mature into signaling endosomes (Füllbrunn et al., 2024), if maturation involves fusion with vacuole-derived nascent signaling endosomal membrane.

Also mysterious is the reason for the heterogeneity within a cell population. It is unclear why only a proportion of cells form signaling endosomes even when Atg18 expression is synchronized (Figure 3B). Whether this difference originates from different cell cycle stages, or different replicative ages, could be examined in the future. Alternatively, because signaling endosome formation requires TORC1 activity (Figure 5), only cells with TORC1 activity above a certain threshold may generate detectable signaling endosomes.

The role of signaling endosomes in intracellular protein trafficking remains a poorly understood aspect of this organelle. Atg18 mediates PI(3,5)P_2_-dependent retrograde trafficking from the vacuolar membrane to canonical endosomes (Dove et al., 2004). Mechanistic details of this process are unknown, in particular whether this trafficking initiates from a specific site of the vacuolar membrane. The role of Atg18 in signaling endosome formation discovered here raises the possibility that signaling endosomes provide the route for retrograde trafficking. If so, the selective migration of vacuolar proteins to newly formed signaling endosomes could serve as a cargo selection process. A fusion event between the vacuole-derived membrane and canonical endosomes might be central for Atg18-mediated retrograde trafficking.

The present study provides our first insight into signaling endosome biogenesis, opening many fundamental questions. Previous studies have identified several protein machineries controlling the abundance and/or the signaling function of signaling endosomes, including the Fab1, HOPS, and ESCRT complexes (Chen et al., 2021)(Gao et al., 2022). A limitation of observing steady-state cell populations however meant that these studies did not discriminate effects on the initial formation, maintenance, and degradation of signaling endosomes. The Atg18 synchronization system we have developed here allows us to dissect each step over time, making it possible to investigate the dynamic life cycle of signaling endosomes.

## Materials and methods

### Yeast strains, plasmids, and growth conditions

*Saccharomyces cerevisiae* yeast strains and plasmids used in this study are listed in Table S1 and S2, respectively. Yeast strains were made prototrophic with indicated plasmids and/or empty vector plasmids, and grown in synthetic complete media without uracil, leucine, and histidine (SC-ULH; SD Broth / 2% Glucose [FORMEDIUM, CSM0210] plus Complete Supplement Mixture Triple Drop-Out - His, -Leu, -Ura [FORMEDIUM DCS0991]). For the *GAL* shut-off experiment, cells were first cultured in synthetic raffinose media and then in synthetic galactose media for 3 hours. After being washed, cells were cultured in SC-ULH media for 3 hours before imaging.

In all experiments, yeast strains were grown to the mid-log phase at 30 °C.

### Fluorescence Microscopy

Images of live fluorescent cells were captured with a VoX spinning disk confocal microscope (Perkin Elmer) equipped with a Flash 4.0V3 camera and a PlanApo VC 60x/1.4 Oil DIC N2 objective.

For time-lapse imaging, live cells were immobilized on a glass bottom dish (ibidi, 81158) coated with 1 mg/ml concanavalin A (Sigma-Aldrich, C2010). After being washed, at t=0, cells were treated with SC-ULH media containing the indicated final concentration of anhydrotetracycline (aTc, Cayman Chemicals, 10009542). Images were obtained with a ZEISS LSM880 confocal microscope (Carl Zeiss) equipped with a PlanApo 63x/1.4 Oil DIC M27 objective and ZEN black software (Carl Zeiss). The temperature was kept at 30 °C throughout the time course. FM4-64 (Invitrogen, T13320) staining was performed as previously described (Hatakeyama et al., 2019) prior to cell immobilization.

Images were processed using Fiji-ImageJ software (National Institutes of Health, http://rsb.info.nih.gov/ij/).

### Western blotting

Yeast cells were treated with 6.7 % trichloroacetic acid (final concentration), pelleted, washed with 70 % ethanol, dissolved in urea buffer (50 mM Tris-HCL [pH8.0], 5 mM EDTA, 6M urea, 1% SDS, Pefabloc and PhosSTOP), and disrupted with glass beads using a FastPrep-24 homogenizer (MP Biomedical). Samples were heated at 65 °C for 10 min in Laemmli SDS sample buffer and subjected to SDS-PAGE and immunoblotting experiments. The following antibodies were used: mouse anti-FLAG (1:5000, Sigma-Aldrich F1804), rabbit anti-Tor1 (1:1000, gifted from T. Maeda), goat anti-mouse IgG-HRP (1:3000, Invitrogen 626520), goat anti-rabbit IgG-HRP (1:3000, Invitrogen A16096).

### Statistical analysis

Two-sided Welch’s *t*-tests were used to calculate *P*-values (unpaired tests for data points in black; paired tests for color-coded data points). The *P*-values with an asterisk were less than the significance levels (α=0.05) corrected by the Holm-Sidak method.

## Abbreviations

TORC1: Target of Rapamycin Complex 1
CROP: Cutting Retromer-on-PROPPIN
PI(3,5)P_2_: phosphatidylinositol 3,5-bisphosphate
PI3P: phosphatidylinositol 3-phosphate

## Acknowledgments

We thank Claudio De Virgilio (University of Fribourg) and Christian Ungermann (Osnabruck University) for yeast strains, plasmids, and for helpful suggestions; Takahiro Shintani (Tohoku University) for plasmids and helpful suggestions; Maya Schuldiner (Weizmann Institute of Science) and Michael Knop (Heidelberg University) for yeast strain libraries; Takeshi Noda (Osaka University), Satoshi Okada (Kyushu University, NBRP plasmid BYP9806), and Joerg Stelling (ETH Zurich, Addgene plasmid FRP2350/2365) for plasmids; Tatsuya Maeda (Hamamatsu University School of Medicine) for the anti-Tor1 antibody; Daniel Paterson, Patryk Marcinkowski, Eri Hirata, and Saran Babooraj (previously University of Aberdeen) for their contribution to the visual screening using YeastRGB; members of the Hatakeyama lab and Chromosome & Cellular Dynamics Section (University of Aberdeen) and Tokai TOR Conference for discussion; Megan Robertson, Colin Ferguson, Arrosan Rajalingam, and Microscopy and Histology Core Facility members (University of Aberdeen) for technical support; Anne Donaldson (University of Aberdeen) for advising on the manuscript. This work was funded by BBSRC (BB/V016334/1 and BB/X018229/1 to RH) and the University of Aberdeen. KM is a recipient of the JASSO scholarship. The authors declare no conflict of interest.

## Author contributions

Conceptualization: KM and RH; Methodology: KM; Investigation: KM; Formal Analysis: KM; Validation: KM; Supervision: RH; Funding acquisition: RH; Visualization: RH; Writing – original draft: KM and RH; Writing – review & editing: RH.

## Tables

**Table S1.**
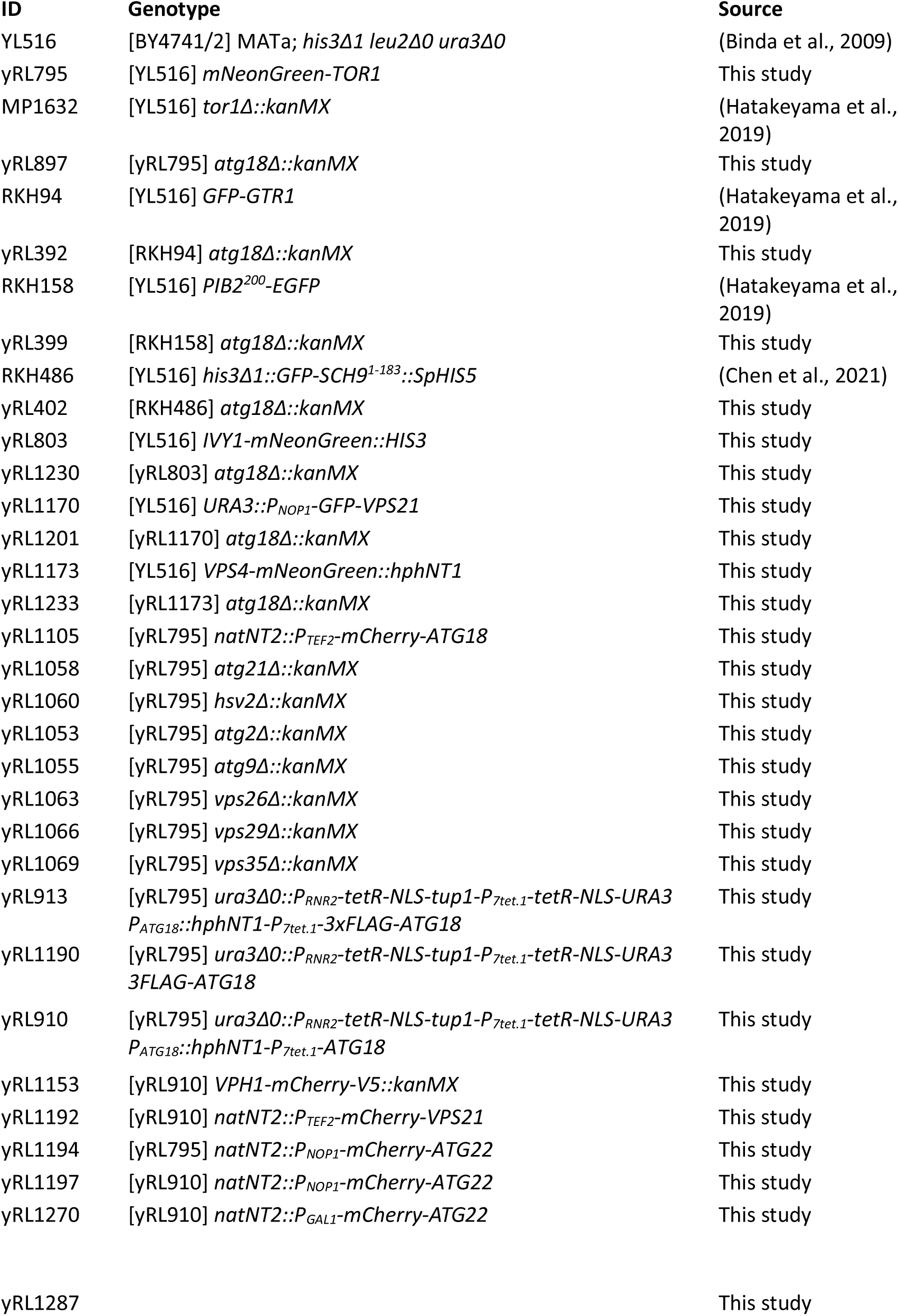
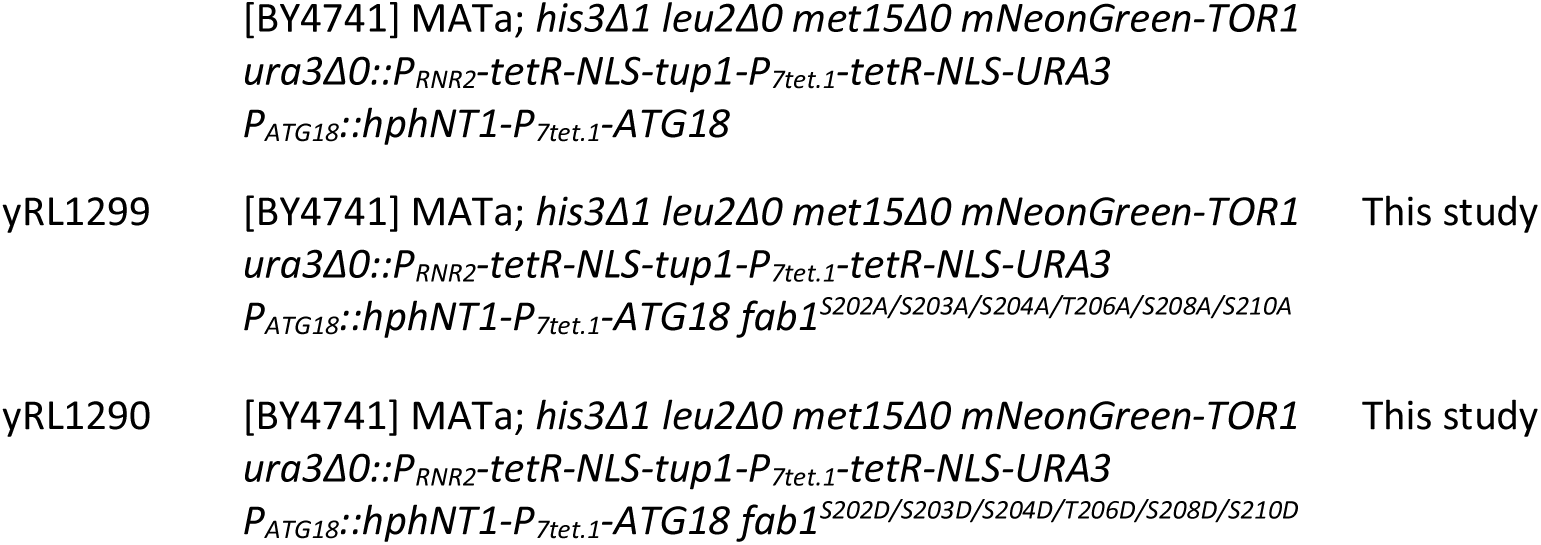
Yeast strains used in this study.

**Table S2.** Plasmids used in this study.

| ID | Name | Description | Source |
| --- | --- | --- | --- |
| pRL127 | pRS413 | CEN/ARS, *HIS3* | (Sikorski & Hieter, 1989) |
| pRL129 | pRS415 | CEN/ARS, *LEU2* | (Sikorski & Hieter, 1989) |
| pRL130 | pRS416 | CEN/ARS, *URA3* | (Sikorski & Hieter, 1989) |
| pRL479 | p1379 | CEN/ARS, *MET15* | (Hatakeyama et al., 2019) |
| pRL910 | pML104i/gTOR1-N | 2μ, *URA3*, *P<sub>GAP</sub>-SpCas9*, *P<sub>SNR52</sub>-gTOR1-N* | (Kira & Noda, 2021) |
| pRL886 | BYP9806 | *CSE4-mNeonGreen-HIS3* | (Okada et al., 2021) |
| pRL913 | pRL913 | *P<sub>TOR1</sub>-mNeonGreen-linker-TOR1<sup>1-426</sup>* | This study |
| pRL942 | FRP2365 | Integrative, *URA3*, *P<sub>RNR2</sub>-tetR-NLS-tup1-P<sub>7tet.1</sub>p-tetR-NLS* | (Azizoğlu et al., 2021) |
| pRL940 | FRP2350 | *hphNT1-P<sub>7tet.1</sub>-3xFLAG* | (Azizoğlu et al., 2021) |
| pRL1007 | pRS416-3FLAG-ATG18 | CEN/ARS, *URA3*, *3xFLAG-ATG18* | This study |
| pRL1022 | pRS416-3FLAG-ATG18T56E | CEN/ARS, *URA3*, *3xFLAG-ATG18<sup>T56E</sup>* | This study |
| pRL1025 | pRS416-3FLAG-ATG18SLoop | CEN/ARS, *URA3*, *3xFLAG-ATG18<sup>Y367K/D369M/K373Y/M375D</sup>* | This study |
| pRL1027 | pRS416-3FLAG-ATG18FGGG | CEN/ARS, *URA3*, *3xFLAG-ATG18<sup>R285G/R286G</sup>* | This study |

